# Comparing advanced interference suppression algorithms for OPM-MEG

**DOI:** 10.64898/2026.09.08.750027

**Authors:** Christoph Pfeiffer, Daniel Lundqvist

## Abstract

On-scalp MEG using optically pumped magnetometers (OPMs) have become more widespread with the recent introduction of the first whole-head systems that offer comparable spatial sampling to conventional MEG systems. For these systems to become truly useful in neuroscience and clinically they need to be able to deal with the kind of interference seen in typical lab and hospital environments – both environmental and participant related (e.g., dental wires). Conventional MEG uses sophisticated interference suppression algorithms to deal with this, but these tend to become instable at lower sensor counts – which makes up the majority of OPM-MEG systems to date. Alternative methods for OPM-MEG are needed. Several alternative algorithms have been proposed but a clear comparison between them is lacking.

Here, we test and compare 5 of the most promising candidates, namely homogenous field correction (HFC), adaptive multipole models (AMM), signal space projection (SSP), signal space separation (SSS) and its iterative implementation (SSS_it_) using simulations and data recorded with a whole-head system. Recorded data includes a participant with dental wire – an especially difficult to deal with source of interference.

We find that all algorithms tested achieve good interference suppression with dual-axis data. With single-axis data, SSS exhibited stability problems and several algorithms showed large signal loss (up to 25%). SSP, temporal SSS and temporal AMM were able to suppress the dental artifact. Finally, we provide recommendations for researchers to determine help decide how to deal with interference in their OPM-MEG studies.

## 1 Introduction

Recording magnetoencephalography (MEG) requires sophisticated magnetic shielding to separate the weak neuromagnetic fields generated by neural activity from the much larger interference fields constantly surrounding us. These interferences include the earth’s magnetic field but can also originate from electronics such as phones and computers, cars, elevators or trains moving close by (Puce & Hämäläinen, 2017). Magnetically shielded rooms (MSRs) use high permeability metal to suppress magnetic fields, lowering them on the inside sufficiently to detect neuromagnetic signals (Cohen, 1968; Hämäläinen et al., 1993). However, even suppressed, interference can significantly surpass the strength of MEG signals with amplitudes in the nanotesla range (Iivanainen et al., 2019). Some interference sources can furthermore not be moved outside the MSR. This includes stimulus (e.g., electric stimulators) and response equipment (e.g., cameras or eye trackers) but also participant related physiological (such as eye and neck muscles, as well as the heart) and artificial sources (such as dental wires/braces, pacemakers, or metal implants) (Hari et al., 2018; Hari & Puce, 2023). For physiological artifacts there are established procedures involving concurrent measurement of eye and heart activity using electrooculogram (EOG) and electrocardiogram (ECG), respectively, to identify and project out relevant interference components (Vigario et al., 2000). Artificial interference sources are more difficult to deal with. While many academic MEG studies can exclude participants with dental wires or pacemakers, clinical recordings as well as studies involving rare patient populations do not have that option. Conventional MEG using cryogenic superconducting quantum interference device (SQUID) sensors, typically employs advanced interference reduction algorithms like signal space separation (SSS) (Taulu et al., 2004) and synthetic higher-order gradiometers (Vrba & Robinson, 2001) to deal with these and other strong interference sources. Such methods require a large number of sensors, however (Holmes et al., 2023). SQUID-MEG typically employs on the order of 300 sensors.

Measuring closer to the participant’s head, optically pumped magnetometers (OPMs) have shown great promise for MEG. Simulations have shown higher sensitivity, more extracted information and higher spatial precision compared to conventional MEG (Boto et al., 2016; Iivanainen et al., 2017; Riaz et al., 2017). First recordings, including in epilepsy patients, using systems with limited sensor count have hinted at the capability of the technology (Feys et al., 2022; Hillebrand et al., 2022). Recently, first whole-head OPM-MEG systems approaching – and in some cases exceeding – 100 channels have become available (Alem et al., 2023; Rea et al., 2022; Schofield et al., 2024; Xu et al., 2025). To get the most out of these systems and make them viable for clinical use, advanced noise suppression methods similar to those used in SQUID-MEG are needed. Several such methods, some specific to OPM-MEG, have been proposed over the last years. While they all provided evidence of their ability to suppress interference, an in-depth investigation into their effects on different types of interference and, importantly, the signal of interest that would allow researchers to make an informed choice about which method to use has been missing.

In this work we test and compare some of the most promising algorithms using realistic simulated, as well as recorded OPM-MEG data. Specifically, we tested: (i) homogenous field correction (HFC), (ii) signal space separation (SSS), (iii) adaptive multipole models (AMM) and (iv) signal space projection (SSP).

### Algorithms

HFC is based on the assumption that magnetic fields from a faraway source can be approximated as spatially homogenous (Tierney et al., 2021). Homogenous fields are therefore assumed to arise from sources outside the sensor array and thus represent interference. The algorithm uses the sensor orientations to compute a projector that projects out homogenous fields. Some implementations of HFC use spherical harmonics expansion which then allows straightforward projecting out of higher spatial orders like the first and second order spatial gradients. We refer to these as HFC here despite the name not being entirely accurate higher orders are used.

SSS is based on the assumption that MEG signals can be separated into those originating from within (representing brain signals) and those originating outside the sensor array (representing interference) (Taulu et al., 2004). Using a spherical harmonics multipole expansion, SSS tries to decompose the measured signals into an inner and outer signal space and then suppress the outer signals by projecting out the outer signal space. To better suppress noise sources, close to the sensor array (e.g., dental wire interference) that leak into the internal space, SSS can be expanded to include a temporal component. Temporal SSS or tSSS tries to identify and remove signal components of the inner space that highly correlate with the residual fields, an indication that they originate in the intermediate space between the inner and outer spaces.

SSS, or its proprietary version MaxFilter (MEGIN Oy, Helsinki, Finland), is regularly used with commercial SQUID-MEG systems but is known to show stability problems in OPM-MEG systems with lower sensor counts (Holmes et al., 2023). Holmes and colleagues thus proposed expanding SSS by using an iterative approach for determining the projector weights (Holmes et al., 2023). The method assumes that the SSS vectors represent the signals in a hierarchical order and iteratively constructs the weights to remove increasingly higher order.

AMM is based on similar assumptions as SSS but tries to construct the inner and outer signal spaces using prolate spheroidal instead of spherical harmonics (Tierney et al., 2024). The idea being that a prolate spheroid better matches realistic head shapes and thus can better encapsulate the entire brain without intersecting the sensor array compared to a sphere. AMM further implements an orthogonal projection such that the filtered signal contains only the inner signal space components orthogonal to the outer signal space as opposed to the oblique projection used in SSS. Like SSS, AMM can be extended with a temporal component by identifying signal components that highly correlate between the inner and residual signals and are therefore assumed to not originate in the inner space.

SSP uses reference data containing noise without the signal of interest such as, for example, an empty room recording without the participant present. The algorithm then extracts principal components from the reference data using principal component analysis (PCA) (Uusitalo & Ilmoniemi, 1997). Assuming these principal components reflect interference sources, a projector can be constructed that projects the principal component(s) out of the data. Assuming the noise is stable between the reference data and the recording, the same projector can be applied to the recording data to suppress the interference.

## 2 Methods

### 2.1 Ethics

We recorded 2 healthy participants with normal hearing and no history of mental illness (left/right-handed=1/1, male/female=1/1, mean age=30). One of the participants had a fixed dental retainer. The study followed the ethical principles for experiments involving human participants laid out by the Declaration of Helsinki and was approved by the Swedish Ethical Review Authority (Dnr: 2023-03283-01).

### 2.2 Recording

The recordings were done inside a 2-layer magnetically shielded room (Ak3B from Vacuumschmelze GmbH, Hanau, Germany) at the Swedish National Facility for Magnetoencephalography (NatMEG) at the Karolinska Institute, Stockholm, Sweden. The same paradigms were recorded in a single visit with whole-head OPM-MEG in single and multi-axis mode using a HEDSCAN system from Fieldline (Fieldline Inc., Boulder, CO, USA). The system contains 136 3rd-generation OPM sensors mounted in an adult Smart Helmet with depth-adjustable slots that automatically detects sensor positions and orientations at the beginning of the recording (Alem et al., 2023). The helmet was fixed to a hight-adjustable MEG-compatible chair (MEGIN Oy) using a custom-made wooden mount. The OPMs used can record a single (Z) or 2+1 axes (hereon called dual-axis), the latter meaning they implement closed-loop operation of all three axes but only use two (Z+Y) for measurement. The sensors are mounted in the helmet such that their Z-axis is oriented radially to the head and the Y-axis tangentially with sensors being alternatingly rotated by 0 or 90 degrees around Z, creating a checkerboard-like pattern. Closed-loop operation extends the dynamic range of the sensors and, when implemented along all three axes, minimizes cross-axis projection errors (Borna et al., 2022). Due to the systems comparatively high dynamic range, no active shielding was used. Electrooculography (EOG, vertical and horizontal) and electrocardiography (ECG) were recorded simultaneous with OPM-MEG using the bio channels of the TRIUX SQUID-MEG system (MEGIN Oy, Helsinki, Finland) housed in the same shielded room – although neither was used in the analysis described here. A computer with a NI data acquisition card (National Instruments, Austin, TX, USA) running Presentation (Neurobehavioral Systems, Berkeley, CA, USA) was used to control stimulation and triggers. MEG triggers were sent to the digital input port of the HEDSCAN system.

A structural MRI was recorded with a 3T SIGNA Premier scanner (GE Healthcare Technologies, Inc., Chicago, IL, USA) during a separate visit at the MR center at the Karolinska Institute. For co-registration, 4 head-position indicator (HPI) coils (MEGIN Oy, Helsinki, Finland) were attached the participant’s head and digitized along with the fiducials and head shape using a Polhemus Fastrak (Polhemus, Colchester, VA, USA). The HPI coils were driven by the HEDSCAN system using a custom adapter box.

In the case of the recording with dental wire, several frontal sensors did not initialize when the participant was in place. This was likely caused by the large, changing field from the artifact since they worked during empty room. As a result, the data with dental artifact only included 107 and 216 channels with single- and dual-axis recording mode, respectively, compared to 129 and 257 channels in the data without dental artifact.

### 2.3 Paradigm

An auditory paradigm was recorded where the participant was presented binaurally with 1kHz tones (300 ms duration). A total of 300 trials were recorded with a variable interstimulus interval (randomized 900, 1400 or 1900 ms) with 200 ms jitter.

Three minutes of empty room data was recorded before the participant entered the MSR with the sensors flush with the inside of the helmet (hereon called “empty room before”), and again after the recording moving the sensors as little as possible (“empty room after”).

### 2.4 Co-registration

The structural MRI was segmented using FreeSurfer (Dale et al., 1999). The resulting cortical sheet was used to create a sourcemodel with 15 684 evenly distributed sources using HCP workbench. A single-shell headmodel was created using Fieldtrip and aligned with the points of the digitized head shape. To complete co-registration of the data the HPI coils were fit to the OPM-MEG data using magnetic dipole fits and the resulting dipoles aligned to the head digitization using an iterative closest points (ICP) algorithm. The resulting transform was then applied to the positions and orientations in the OPM sensor definition.

### 2.5 Preprocessing

The data was high pass-filtered at 1 Hz and lowpass-filtered at 70 Hz using 4^th^-order Butterworth IIR filters. Bad channels were automatically identified and excluded based on i) low correlation (<0.6) with closest neighbours, ii) flat data segments, iii) missing sensor location, and iv) spectral outliers (>3 standard deviations above the mean at > 50% of the spectrum).

### 2.6 Interference reduction algorithms & Analysis

After removing bad channels, the data was processed using the different spatiotemporal algorithms. Where possible algorithms were implemented using existing functions. For HFC, AMM and SSS the Fieldtrip functions *ft_denoise_hfc*, *ft_denoise_amm* and *ft_denoise_sss* were used. For the iterative SSS method introduced by Holmes and colleagues, *ft_denoise_sss* was modified, adding the function for iterative selection of the weights published along with their paper (Holmes et al., 2023). HFC takes the order as an input, SSS, SSS^it^ and AMM the inner L_in_ and outer order L_out_. Regular SSS and AMM further have a temporal correlation threshold thr with thr<1 leading to the temporal implementation of the algorithms being used. SSP was implemented by computing the principal components from a reference noise recording, e.g., empty room, using the MATLAB function *pca* with the number of principal components n_PC_ as input variable. The projector P to project the principal components defined by the coefficient matrix V_PC_ out of the data was then computed as:

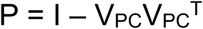

Where I denotes the identity matrix. The projector was then applied to the data trials. To ensure consistency the sensors and their order was matched between the reference and recording data. Three different options for the reference data were tested: empty room before, empty room after, and the pre-stimulus period (-100 to 0 milliseconds) of the auditory experiment. The different versions are hereon referred to as SSP_pre_, SSP_post_ and SSP_data_, respectively.

The following parameters (and combinations thereof) were tested:

- HFC: order = 1,2,3
- SSS, SSS_it_ and AMM:
  - L_in_ = 8, 9, 10
  - `o` L_out_ = 1, 2, 3
  - thr = 0.8, 0.9, 0.99, 1 (only SSS and AMM)
- SSP: n_PC_ = 1, 2, 3, 4, 5, 6, 7, 8, 9, 10

In the case of SSS,SSS_it_ and AMM, only parameter combinations resulting in a total number of basis vectors smaller than the number of channels n_ch_ were tested (Holmes et al., 2023):

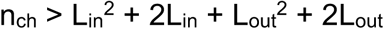

### 2.7 Data analysis

After applying the interference suppression algorithms, the cleaned data was segmented into trails -100 to 500 ms, notch-filtered (50 and 60 Hz using discrete Fourier transform filters), demeaned (baseline = pre-stimulus period) and averaged over trials. Peak latencies and amplitudes were extracted automatically as the maximum absolute fields within a window (±10 ms) around the main response component, namely the M100. The signal-to-noise ratio (SNR) of the peak M100 activation was calculated as the peak amplitude of the averaged response *x̅* divided by the standard error over trials at the same channel and latency as the peak amplitude.

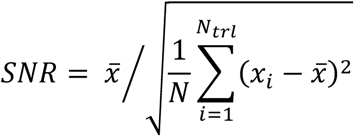

Two symmetric (in X) current dipoles were fitted to the peak latency identified from the averaged responses and locations and residual variances (RV) of the fit extracted.

Finally, the evoked data was source reconstructed using minimum norm estimates (MNE), with fixed source orientation, source covariance scaling and regularization parameter lambda = 3. A noise covariance matrix calculated from the empty room before data was used for whitening. The peak latency and location, defined by the maximum source power in the response component window, along with the extend of the peak activation of the MNE reconstruction were then extracted. The extend was defined as the full area at half maximum (FAHM), i.e., the sum over the source area with an amplitude greater or equal to half the maximum source activation. The FAHM was calculated separately per hemisphere and subsequently averaged.

Since the ground truth is not known in case of the recorded data, the methods are evaluated primarily based on sensor level SNR, dipole fit RV and MNE reconstruction FAHM. The latter two are based on the knowledge from literature that the M100 activates small bilateral patches in the primary auditory cortex in the superior temporal gyrus that can be approximated well by dipoles (Hari et al., 2018). Clean data should therefore lead to low RV when fitting dipoles and a small activation area – and hence a small FAHM, when doing a MNE source reconstruction.

### 2.8 Simulated data

Using the headmodel, sourcemodel and sensor definition from the recording without dental wire, we simulated a recording with different brain signals and noise sources. Three versions were simulated representing a system with sensors at 130 locations recording singe-axis (130 channels, radial only), dual-axis (260 channels, radial and one tangential axis), and triaxial (390 channels, radial and both tangential axes). The sensor signals were calculated as the sum of i) sensor noise, ii) signals from noise sources, and iii) signals from brain sources.

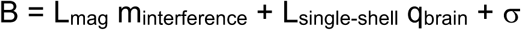

where L_mag_ and L_single-shell_ denote lead fields for magnetic dipoles in infinite vacuum and electric dipoles in a single-shell headmodel, respectively, **m** and **q** magnetic and electric dipoles and **σ** sensor noise.

#### Sensor noise

Sensor noise was simulated for each OPM sensor as the sum of randomly generated pink and white noise generated using MATLABs dsp.ColoredNoise function. The amplitudes of white (10.9 fT/Hz^1/2^) and pink noise (10.9 fT/Hz^1/2^ at 3 Hz) were chosen to match the average sensor noise in the empty room recording.

#### Interference sources

Interference sources were simulated as magnetic dipoles orientated towards the OPM helmet to maximize the fields they generate at the sensor array. Interference sources were positioned at (i) at 30 cm inferior and anterior to simulate a dental source (for example a dental wire), (ii) 50 cm inferior to simulate a cardiac source (for example, a pacemaker), (iii) 1 (right) and (iv) 2 m (anterior) to simulate sources inside the MSR (for example, stimulation equipment), and (v) 3 (right) and (vi) 5 m (anterior) to simulate faraway sources. The sources were activated with sinusoidal signals at 19, 31, 43, 59, 71, and 83 Hz, respectively, and random phase. The amplitude of the dipoles was scaled such that all sources generated peak signal amplitudes of 141 pT at the sensor array. With a 10.9 fT/Hz^1/2^ white noise floor, this means realistically noise suppression for each source is limited to around 80 dB.

#### Brain sources

To test the effect of the algorithms on brain signals, 20 strongly coupled sources spread throughout the brain were selected (see figure 1, left). A separate trial was generated for each brain source. They were each activated with a sinusoidal signal at 10 Hz scaled to generate a peak signal amplitude of 10 pT.

**Figure 1:**
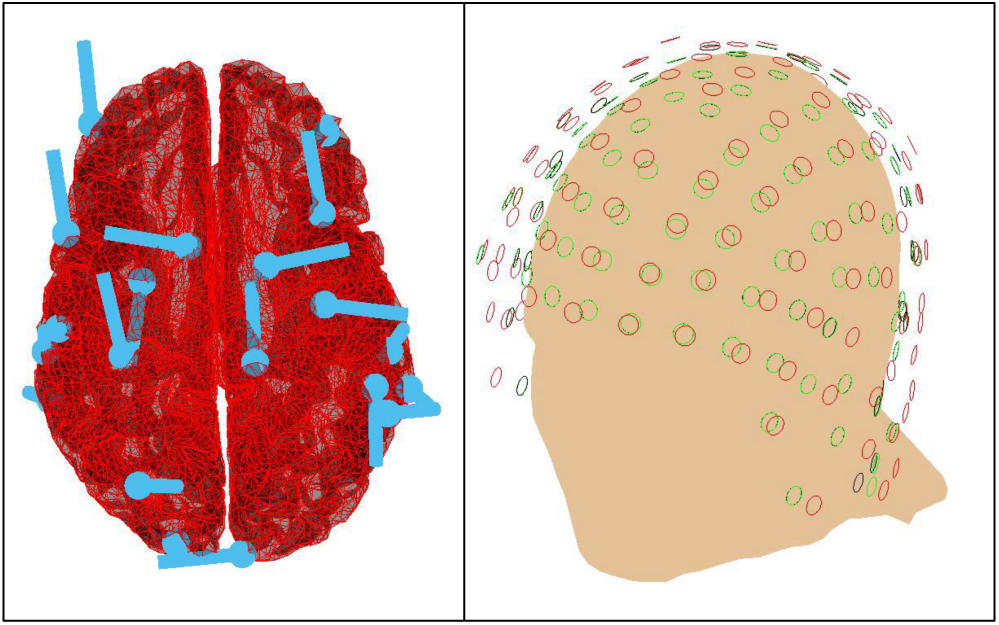
Left: Dipole sources (light blue) in the brain (red) used for simulating signals of interest. Right: Sensor arrays during the before (green), during (black circles) and after the recording (red circles).

#### Empty room data

To simulate an empty room recording for the SSP algorithm, we generated data using the same environmental interference sources (iii-vi) but without the participant related interference sources (i and ii) and brain signals. Three empty room datasets were simulated, using the sensor arrays from the empty room before and after recordings, as well as, from the experiment itself. The sensor array used during the recording represents the ideal case where sensor positions and orientations do not change between the reference data and the data to clean, whereas the array from empty room before represents the largest change. As seen in figure 1 (right), the array from empty room after shows only minimal changes compared to the one used in the experiment.

#### Resting state data

In event related experiments, SSP can also be done using resting state or pre-stimulus data as reference data with the assumption that it contains both environmental and participant-related but task-unrelated interference. To simulate resting state data, a recording similar to the empty room data without brain signals but with all 6 interference sources present was generated.

## 3 Results

### 3.1 Simulated data - single-axis

SSS became instable at all the parameter combinations tested when used on single-axis data. The other spatial algorithms, HFC, SSS_it_ and AMM, showed a similar trend in interference suppression trend with faraway interference sources being suppressed stronger than close ones, and higher order/L_out_ leading to an increase in suppression (see figure 2, top row). At higher order/L_out_, SSS_it_ showed a slightly lower average suppression of environmental interference (56 dB compared to 63 dB with HFC and AMM at order/L_out_ = 3) but at lower signal loss (2% compared to 19% and 23% for HFC and AMM at order/L_out_ = 3, respectively) compared to HFC and AMM. Unlike SSS_it_, both HFC and AMM showed a similar trend of increasing signal loss with increasing order/L_out_ (see figure 2, bottom row).

**Figure 2:**
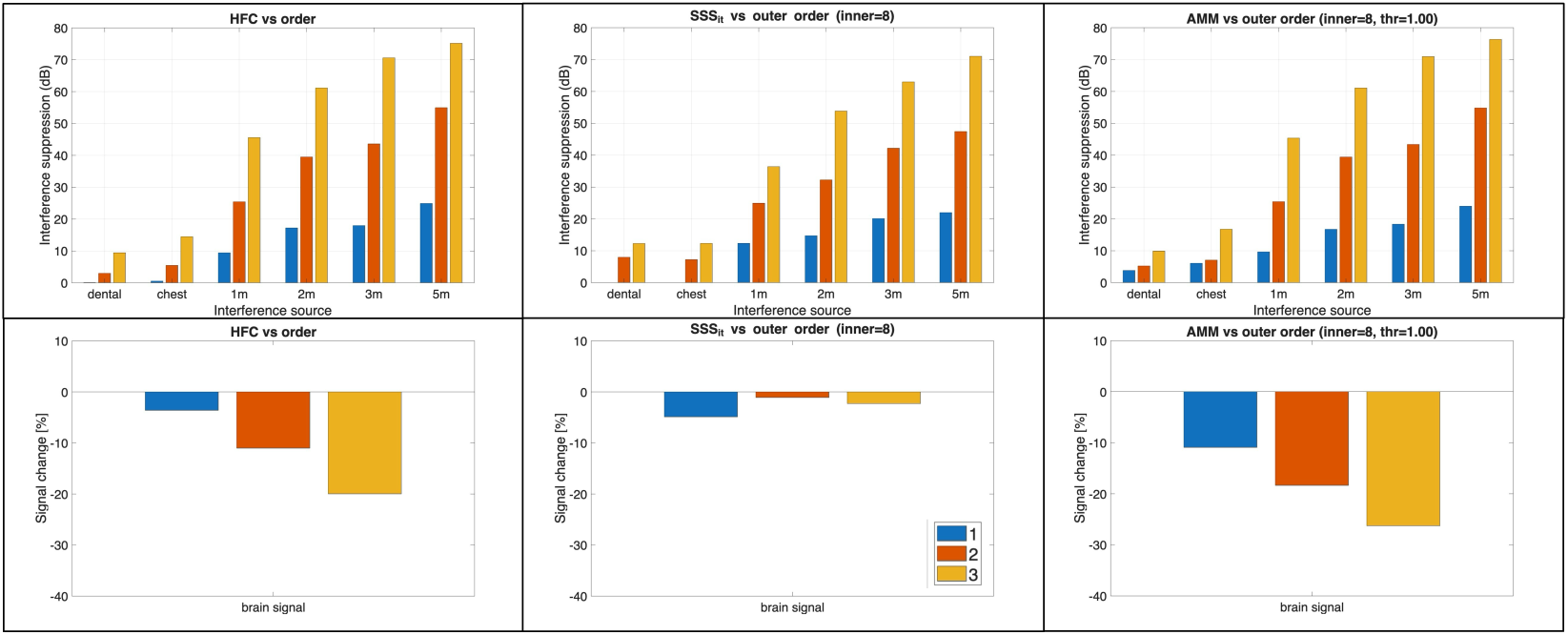
Interference suppression (top) and signal loss (bottom) for HFC (left), iterative SSS (middle), and AMM (right) as a function of order/L_out_ using single-axis data.

L_in_ showed little effect on the noise suppression in SSS_it_ and AMM using single-axis data (see figure 3, top row). AMM and SSS_it_ both exhibited increasing signal strength with increasing L_in_ (see figure 3, bottom row). SSS_it_ overshot, showing a positive signal change at Lin ≥ 9.

**Figure 3:**
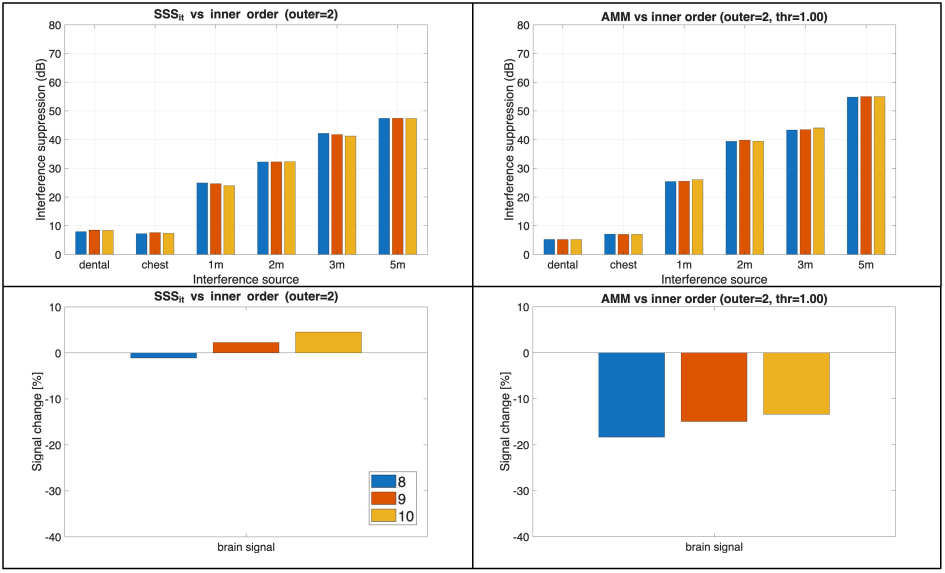
Interference suppression (top) and signal loss (bottom) as a function of L_in_ for iterative SSS (left) and AMM (right) using single-axis data.

SSP using empty room data showed a plateau of the interference suppression at n_PC_ ≥ 4, indicating the first four principal components may have captured all four environmental interference sources (see figure 4, top row). The peak suppression differs between the different empty room data with ideal empty room performing best, followed by empty room after and empty room before, with average environmental interference suppression of 75, 47, and 40 dB, respectively. None of the empty rooms showed much effect on the dental and chest interreference sources, which were not present in the empty room data. SSP using resting state data as reference, on the other hand, reached similar peak suppression as with ideal empty room, including for the dental and chest sources, but only plateaued at n_PC_ ≥ 6 (see figure 4, top right). The later plateau makes sense considering there are two more interference sources present in the reference data. All SSP implementations showed a similar trend of the signal loss increasing with increasing n_PC_ (figure 4, bottom row). Overall, the signal loss remained low with a maximum of 11% at n_PC_ = 10 and ∼6% at the ideal n_PC_ (i.e., the start of the plateau).

**Figure 4:**
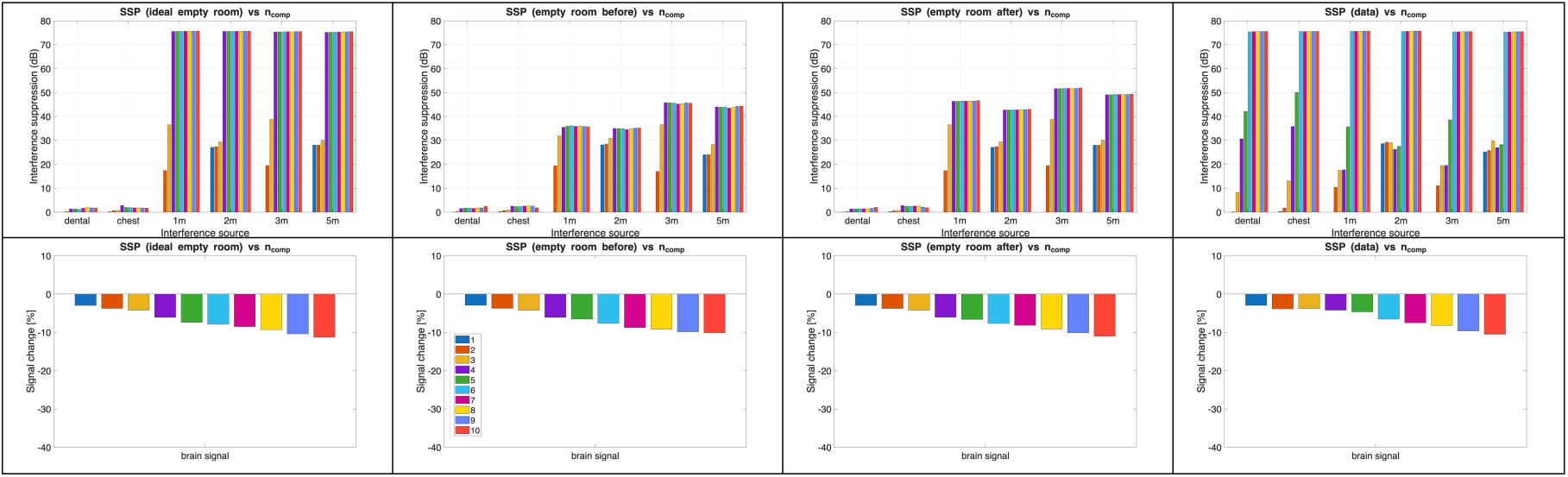
Interference suppression (top) and signal loss (bottom) with SSP using ideal empty room (left), empty room before (middle), and empty room after (right) data as reference for single-axis data.

### 3.2 Simulated data – multi-axis

Regular SSS remained stable with dual and triaxial data. The spatial algorithms showed similar interference suppression with multi-axis data compared to single-axis data. All showed the same trends of faraway interference sources being stronger suppressed and higher order/L_out_ increasing suppression (see figure 5, top row). HFC, and AMM to a lesser extent, showed significantly less (up to 17 and 10%, respectively) signal loss compared to single axis data. Both showed higher signal loss with increasing order/L_out_. SSS_it_ showed an overall increase in signals loss compared to single axis data. Like SSS, the signal loss was not affected by L_out_.

**Figure 5:**
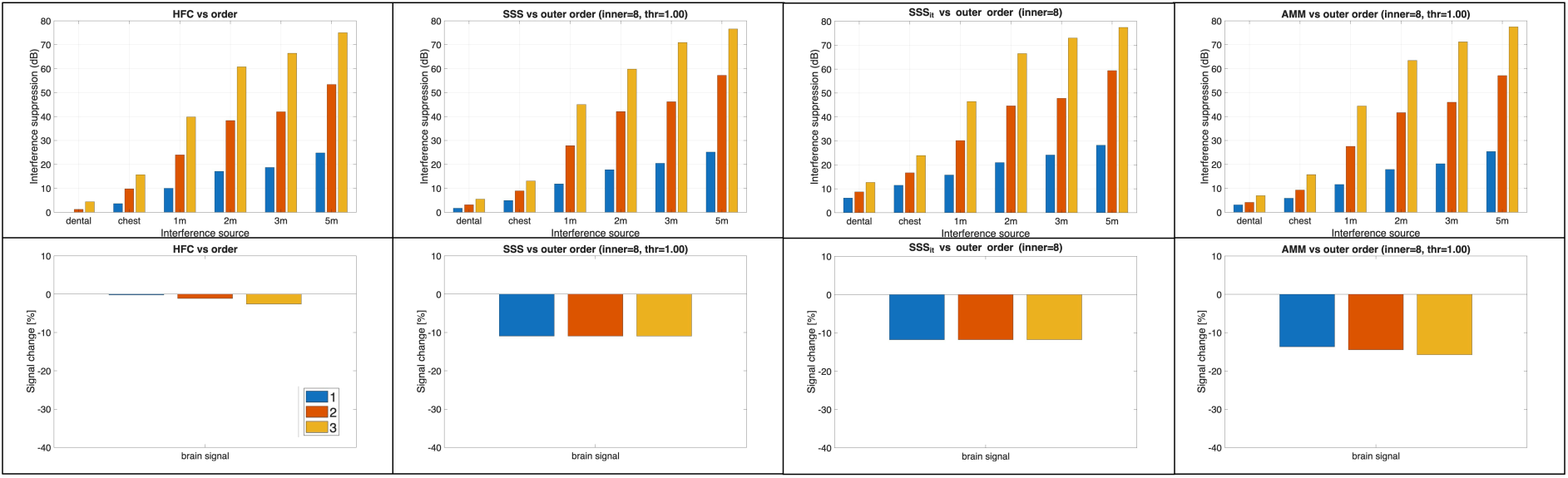
Interference suppression (top) and signal loss (bottom) for HFC (left), SSS (center-left), iterative SSS (center-right), and AMM (right) as a function of order/L_out_ using dual-axis data.

SSS, SSS_it_ and AMM all showed a minor reduction (2,1 and 4 dB, respectively) in interference suppression with increasing L_in_ (see figure 6, top row). Simultaneously the signal loss dropped by 5-7% with increasing Lin (figure 6, bottom row).

**Figure 6:**
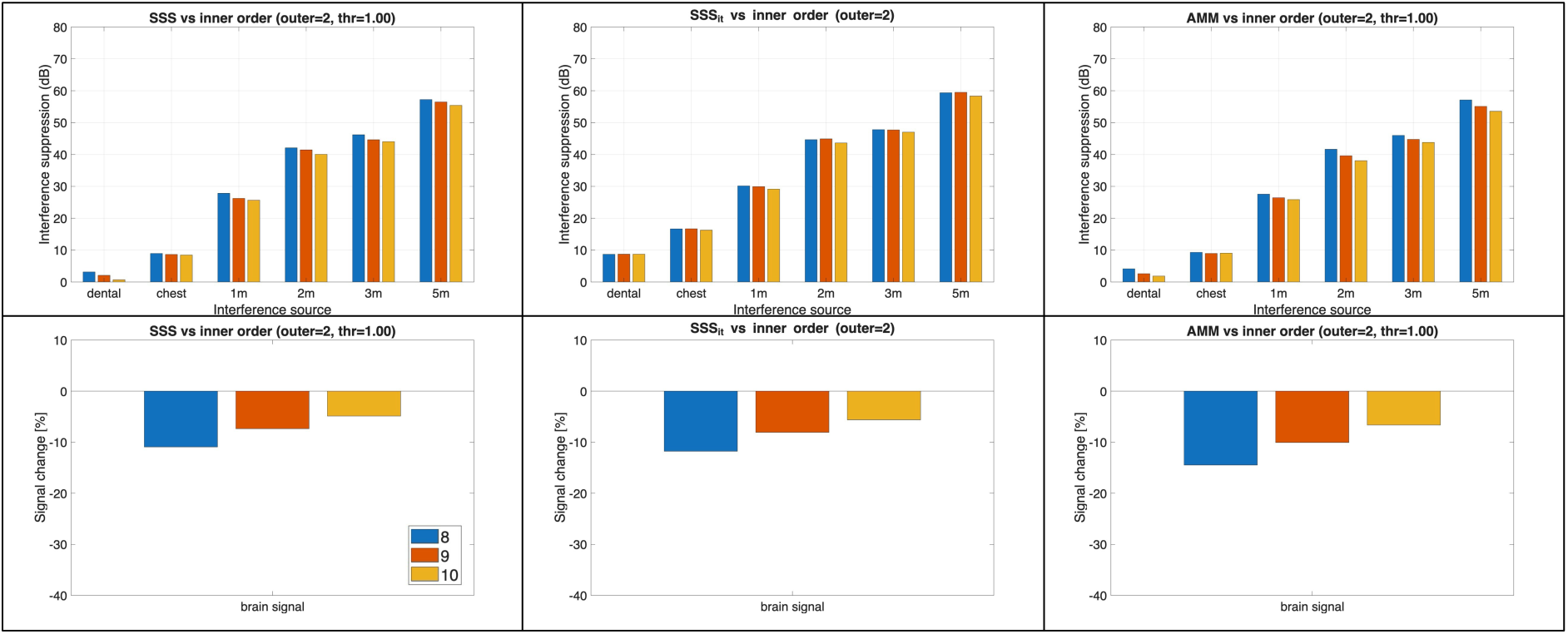
Interference suppression (top) and signal loss (bottom) as a function of L_in_ for SSS (left), iterative SSS (middle) and AMM (right) using dual-axis data.

Interference suppression with SSP showed similar trends with multi axis data compared to single axis data (see figure 7, top row). Suppression plateaued at n_PC_ = 4 and n_PC_ = 6 with empty room and resting state data as reference, respectively, and showed the same drop in peak suppression when going from ideal empty room to empty room after and empty room before. Signal loss was even lower compared to single axis data with a maximum of 4% and less than 1% at ideal n_PC_.

**Figure 7:**
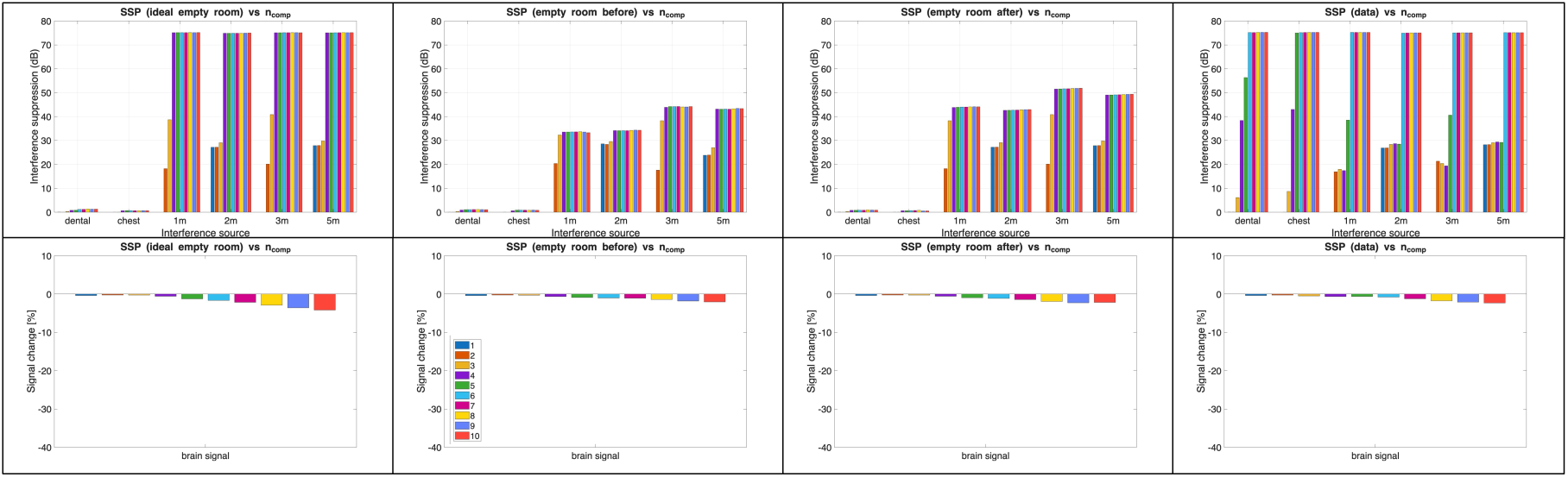
Interference suppression (top) and signal loss (bottom) with SSP using ideal empty room (left), empty room before (middle), and empty room after (right) data as reference for single-axis data.

HFC and SSP showed no significant difference in interference suppression or signal change between dual and triaxial data. SSS_it_ showed a 2-3 dB increase in interference suppression with triaxial data for all parameters tested. SSS, SSS_it_ and AMM three methods also showed a minor increase (∼1.5%) in signal loss with triaxial data.

### 3.3 Recorded data

Most of the algorithms were able to significantly reduce interference. HFC, AMM and SSP increased the sensor-level SNR by 85-92%, with SSP_pre_ performing best. Similarly, the dipole fit RV was reduced by 58-72%, while MNE FAHM showed only little improvement (1-16% reduction).

SSS became instable at L_in_ ≥ 9 leading to an increase in noise. In the best case, SNR only increased by 33% - less than half of most of the other methods. Iterative SSS similarly showed a much lower increase in SNR (+35%) than most of the other algorithms. At L_out_ = 3, a large signal drop (63%) was observed with SSS_it_.

HFC and AMM showed large signal drops (28-31%) at order/L_out_ ≥ 2. In addition to these signal drops, the emergence of “oscillatory”-like spatial patterns in the topography was observed at order/L_out_ > 1 (see figure 8). The spatial frequency of these “oscillations” appears to be increasing with increasing order/L_out_, starting with what looks like a first order gradient (left to right) at order/L_out_ = 2 (figure 8b and 8e), followed by spatially periodic activations between the left and right dipolar patterns of the (bilateral) M100 at order/L_out_ = 3 (figure 8c and 8f).

**Figure 8:**
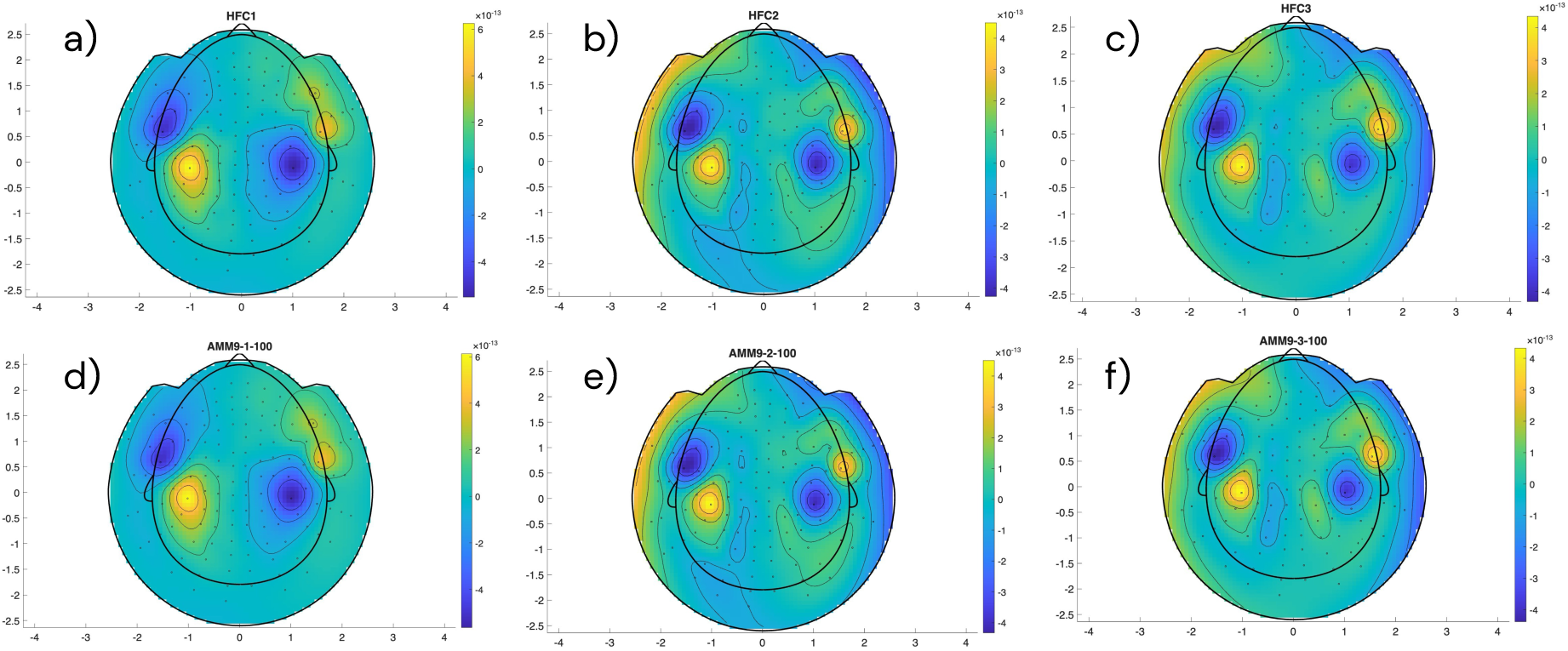
Topographic plots of the M100 after interference suppression with HFC a-c) and AMM (d-f) showing emergence of spatial patterns at increasing order/L_out_ in single-axis data.

As expected from the simulations, SSP using empty room data showed minimal signal drop leading to the highest SNRs of all algorithms. Only minor differences were seen between SSP_pre_ and SSP_post_ indicating both were able to capture the main interference sources effectively. Both showed drops in signal (13 and 10 % for SSP_pre_ and SSP_post_, respectively) at n_PC_ ≥4. SSP_data_ showed a continuous drop in signal from n_PC_ = 4 down to 62% at n_PC_ ≥ 8, leading to a drop in SNR.

The multi-axis recording showed more noise than the single-axis data. Since this increase in noise was not reflected in the empty room data, it is likely the result of stronger interference being present at the time of recording. All methods including SSS and SSS_it_ provided good interference suppression achieving comparable SNR increases (78-112%) as with the single-axis data.

In agreement with the simulations, using dual-axis data greatly improved the signal loss seen with some of the algorithms. While HFC and AMM still showed small signal drops at order/L_out_ ≥ 2, these were much weaker (<8%) compared to the ones observed with single-axis data. They also did not show the oscillatory spatial patterns seen with single-axis data, making the use of higher orders more viable.

With the larger number of channels, SSS and SSS_it_ were stable when used with multi-axis data, achieving, along with AMM, some of the highest SNR and lowest dipole fit RV of all the algorithms. MNE source reconstruction, on the other hand, worsened showing an increase in FAHM for the algorithms compared to no interference suppression. AMM, SSS and SSS_it_ all achieved the highest SNR and lowest RV at the lowest L_in_ tested (L_in_ = 8), while the lowest FAHM was reached at the highest L_in_ (L_in_ = 10). Temporal SSS and AMM both remained stable and led to a minor increase in SNR at L_in_=8, L_out_=2 with correlation threshold ≤0.95. Generally, the effects were minor, as expected, given that there was no major source of interference close to the sensor array.

SSP_pre_ and SSP_post_ showed the lowest improvement in SNR and RV of the algorithms tested. As with single-axis data, the algorithm performed similar with either of the empty room data. SSP_data_ showed a large drop in signal (66% at n_PC_ = 6 down to 83% at n_PC_ = 10) leading to a drop in SNR.

Both, with single and dual-axis data, AMM and SSS performed best with thr < 1, indicating some interference that bled into the intermediate space was present in the data.

Table 1 summarizes the best sensor level SNR, dipole fit RV and MNE source reconstruction FAHM for the different algorithms with single and dual-axis data for the data recorded without dental artifact.

**Table 1:** Best M100 peak sensor level SNR, dipole fit RV and MNE FAHM for each algorithm with single and dual-axis data recorded without dental artifact.

|  |  | None | SSP<sub>data</sub> | AMM | HFC | SSS | SSS<sub>it</sub> | SSP<sub>pre</sub> | SSP<sub>post</sub> |
| --- | --- | --- | --- | --- | --- | --- | --- | --- | --- |
| Singleaxis | SNR | 0.56 | 1.05 | 1.08 | 1.05 | 0.81 | 0.76 | 1.07 | 1.09 |
|  | RV [%] | 16.4 | 6.4 | 4.8 | 6.3 | 7.0 | 7.0 | 7.0 | 6.9 |
|  | FAHM [cm<sup>2</sup>] | 10.4 | 8.7 | 9.6 | 9.2 | 9.5 | 9.7 | 10.3 | 10.2 |
| Multi-axis | SNR | 0.49 | 0.93 | 1.03 | 0.91 | 0.95 | 0.95 | 0.87 | 0.87 |
|  | RV [%] | 26.2 | 11.8 | 9.8 | 11.0 | 10.2 | 9.3 | 14.8 | 13.7 |
|  | FAHM [cm<sup>2</sup>] | 15.9 | 13.7 | 19.3 | 15.6 | 18.6 | 19.5 | 15.8 | 15.9 |

<sup>1</sup>Bad reconstruction/localization

The data with dental artifact showed very strong interference with no distinguishable M100 at sensor level. Dipole fits resulted in sources in the frontal lobe along the midline, likely representing primarily the dental wire artifact. MNE source reconstruction similarly showed a strong, frontal activation.

Most of the algorithms were unable to effectively suppress the dental artifact. With single-axis data, only tAMM (at L_in_ = 8 and thr ≤ 0.9) and SSP_data_ managed to recover a clear, although noisy in the case of tAMM, M100 on the sensor level, as well as reasonable dipole source locations (bilateral superior temporal region). All other methods resulted in incorrect dipole locations. MNE source reconstructions showed M100 activations with prominent peaks in the right temporal lobe with SSP_data_, tAMM, and tSSS (L_in_ = 8 and thr ≤ 0.9).

With dual-axis data, tSSS and tAMM both showed clear M100 activations at sensor and source level for thr ≤ 0.99 (see figure 9), with best SNR at thr ≤ 0.9. As with singe-axis data, tAMM/tSSS performed better at low L_in_. Higher inner orders may result in more of the dental signal being included in the inner signal space, making it less likely for them to be removed.

**Figure 9:**
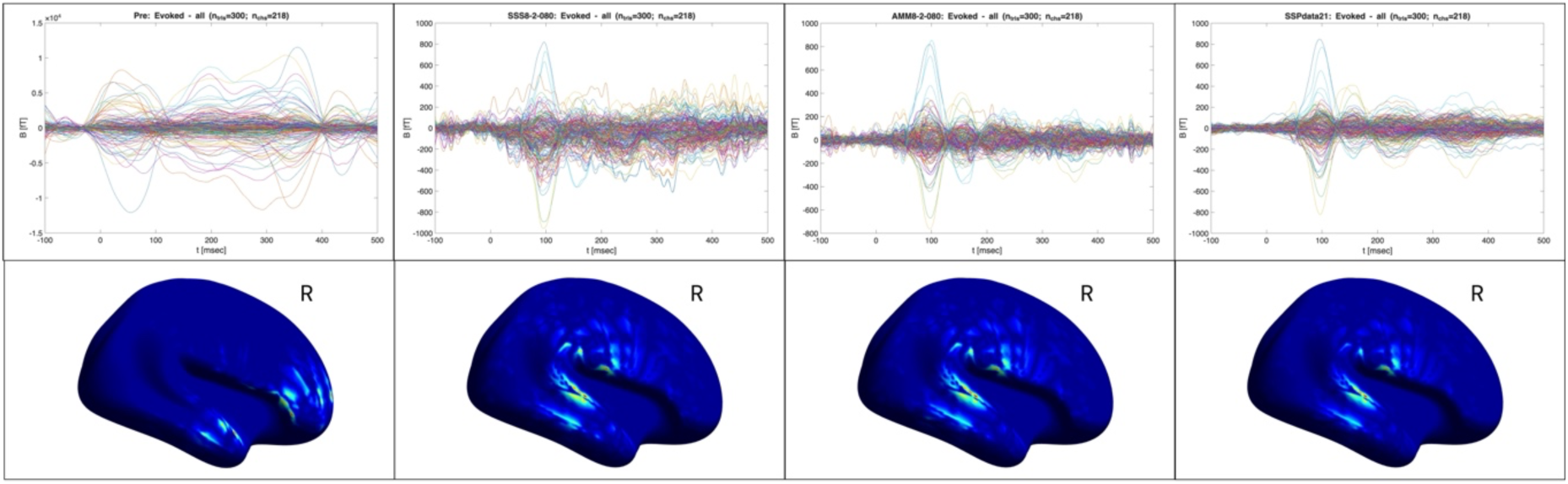
Butterfly (top) and MNE source reconstructions of the M100 (bottom) using the dual-axis dental artifact data without interference rejection (left column), tSSS (center left column), AMM (center right column) and SSP_data_ (right column).

SSP_data_ achieved higher suppression of the dental artifact than any of the other algorithms. Since the SNR did not flatten off at n_PC_ = 10 we tested up to 25 principal components for the dental data and found the highest SNR was achieved with n_PC_ = 15 and 21 with single- and multi axis data. As in the data without dental wire, SSP_data_ exhibited large signal drops in n_PC_ > 19 and n_PC_ > 22 with single- and dual-axis data, respectively, indicating later principal components may include signal of interest.

HFC and SSS_it_, which do not account for temporally varying interference, as well as SSP_pre_ and SSP_post_ which, unlike SSP_data_, did not include the dental artifact in the reference data, showed no noticable improvements in data quality.

Interestingly, all algorithms showed a strong right dominance of the M100, with the left hemisphere activation only barely noticeable (see figure 10). Considering that later components do show activity in the left hemisphere, this may be the result of imprecise positioning of the speakers during the experiment which led to unbalanced sound presentation.

**Figure 10:**
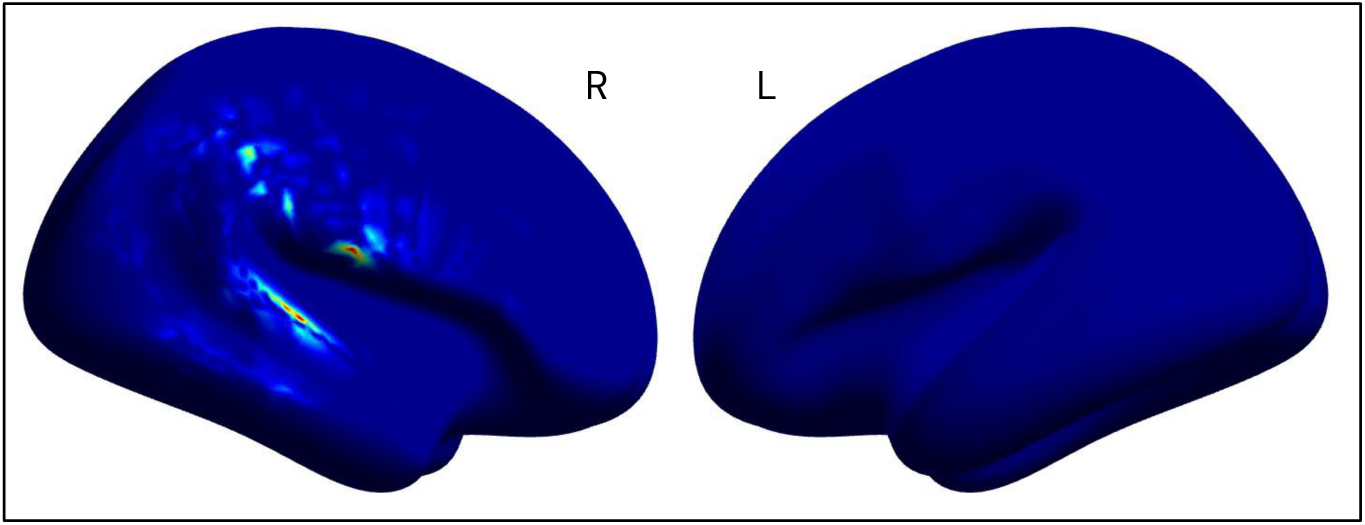
MNE source reconstruction of single-axis dental artifact data without interference rejection using noise covariance matrix calculated from pre-stimulus data.

Table 2 summarizes the best sensor level SNR, dipole fit RV and MNE source reconstruction FAHM for the different algorithms with single and dual-axis data for the data recorded with dental artifact.

**Table 2:**
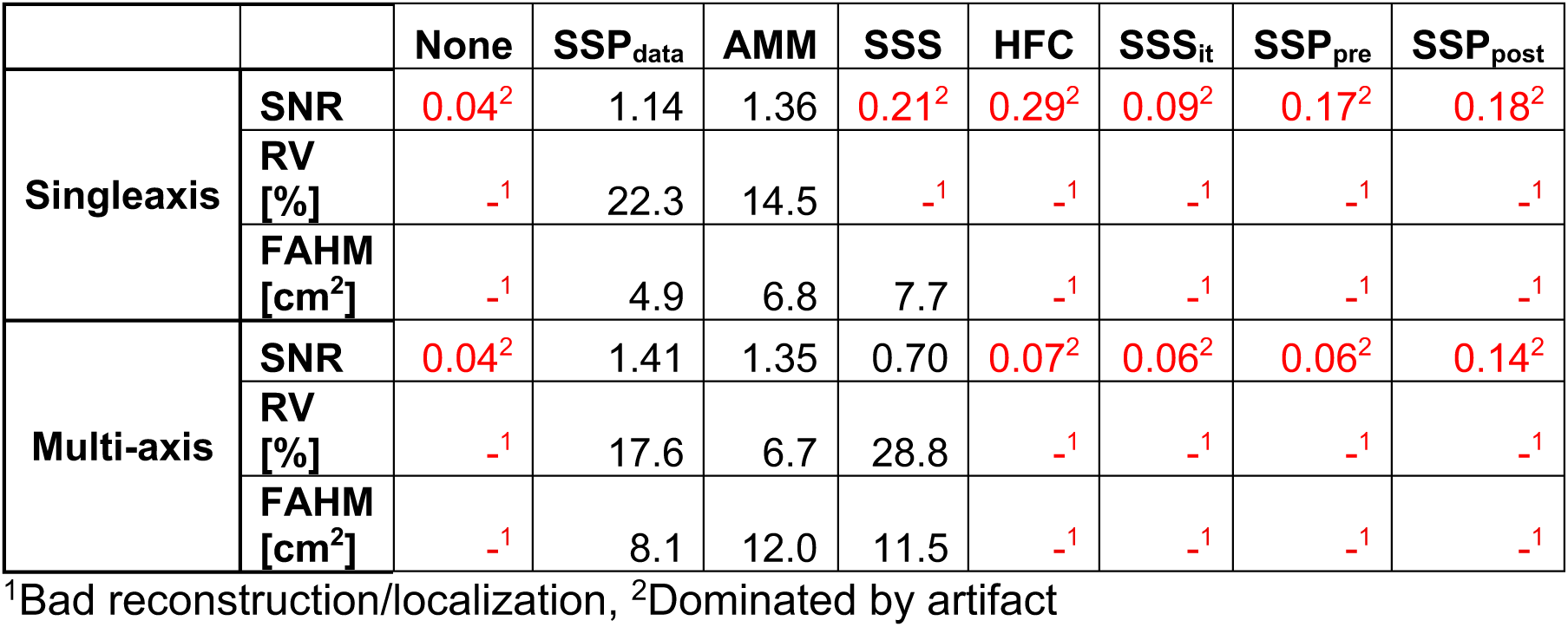
Best M100 peak sensor level SNR, dipole fit RV and MNE FAHM for each algorithm with single and dual-axis data recorded with dental artifact.

We also tested using a noise covariance matrix calculated from the pre-stimulus period, as opposed to empty room data. Using a noise covariance matrix from pre-stimulus data led to successful MNE source reconstruction in almost all cases – including without interference suppression (see figure 10). Only SSS_it_ at some parameters showed incorrect activations. The algorithms show little (positive) effect on the reconstruction in this case, indicating the source reconstruction in itself provides a certain interference suppression.

## 4 Discussion

While it is difficult to compare performance of the algorithms across different OPM systems, our results agree reasonably well with findings from other groups. Tierney and colleagues showed suppression of environmental interference of up to 20 dB for 1^st^-order HFC using a system with 31 dual-axis sensors (Tierney et al., 2021). The fact that we saw similar suppression of faraway interference sources in simulations with 2-6 times as many channels indicates that a few sensors are enough to sufficiently capture the homogenous field for projecting it out of the data. Adding additional channels primarily helps minimizing signal loss. In another study, the same group showed suppression of environmental interference of up to 40 dB with AMM and instability with tSSS using a system with 64 dual-axis sensors (Tierney et al., 2024). Both, AMM and tSSS results fit with our results for data recorded with 129 single-axis sensors. In simulations, as well as in the recorded data without dental artifact, SSP showed the highest SNR at n_PC_ = 4, comparable to the ideal threshold of 3 components for auditory evoked activity recorded with 26 sensors found by Zhao and colleagues using their automated algorithm (Zhao et al., 2024). The low number of components likely reflects typical environmental interference sources such as vibrations of the shielded room and fluctuations in the homogenous field. For somatosensory evoked activity, they found a higher ideal number of components (Zhao et al., 2024). The additional components may reflect the additional interference caused by the electric stimulator. Here, we similarly found the optimal SNR at higher number of components (n_PC_ =15 and 21 for single- and dual-axis data, respectively) when the recording contained the additional dental interference source.

AMM achieved the best performance of the SSS-type algorithms (AMM, SSS and iterative SSS) combining good interference suppression with stability and relatively low signal loss but depends strongly on the parameter selection. SSS and iterative SSS achieved good interference suppression with multi-axis data, but SSS struggled with stability when applied to single-axis data. HFC and SSP performed well in all metrics, both for single- and multi-axis data. Especially with single-axis data, SSP achieved by far the lowest suppression of brain signals of all the methods tested.

Suppression of brain signals can have a significant impact on analysis. This should be kept in mind when analysing sensor-level amplitudes – especially when comparing between different data that are processed differently like studies comparing OPM- and SQUID-MEG.

> *Recommendation 1:* With single-axis data, use SSP or low order/L_out_ for HFC, AMM and SSS_it_.

With single-axis data, HFC and AMM further showed the emergence of spatial patterns at higher orders/L_in_. Considering they do not appear in the multi-axis data, these patterns are likely to be artificial with no underlying neural activation.

Presumably they represent spatial filtering artifacts of the sharp, bilateral activation similar to the ringing caused by sharp changes seen in temporal filters.

Adding a second measurement axis significantly improved signal loss for HFC, AMM and SSP and resulted in SSS becoming stable. Interestingly, adding a third axis did not show significant improvements to noise suppression in simulations compared to two axes. A clear improvement in signal loss for adding the third axis compared to two was only observed with SSS.

> *Recommendation 2:* If possible, record multi-axis data.

SSP showed a drop in noise suppression when using reference data with the sensors in different locations than during the participant recording. While the difference was small in the recorded data, the simulations show an increase in noise suppression of ∼10 dB with SSP_post_ compared to SSP_pre_ where the sensors were further from their positions during the participant recording.

> *Recommendation 3:* If possible, record empty room after with minimal sensor movement.

As seen in the simulation results, SSP only suppresses interference sources that are also present in the reference data. Interference sources related to stimulus, for example, artifacts caused by valves used to control artificial muscles or membranes, should also be present in the reference data if possible.

> *Recommendation 4:* If the stimulation causes artifacts, run it while recording empty room data.

Dental artifacts proved difficult to clean. Of the algorithms tested only tSSS, tAMM and SSP_data_ were able to recover brain activity, the former doing so effectively only with dual-axis data. Temporal SSS and AMM showed best performance at low correlations thresholds (thr ≤ 0.9). A low correlation threshold makes sense given the low SNR in the data. SNR is known to strongly affect the correlation between inner and intermediate spaces and thus the ideal correlation limit (Tierney et al., 2024). SSP_data_ surpassed both methods, especially with single-axis data. However, this method should be used carefully, as it can be difficult to distinguish artefactual and brain components. Removing too many components, can result in the signal of interest being suppressed or distorted. An automated algorithm for selecting the number of components to remove as the one proposed by Zhao and colleagues could help selecting a suitable threshold (Zhao et al., 2024).

> *Recommendation 5:* SSP_data_ and (for multi-axis data) tAMM and tSSS can be used to suppress dental artifacts. Carefully tune the correlation threshold/number of principal components.

Using the pre-stimulus period as reference for calculating the principal components as done here should generally be done with caution. If, due to a short ISI, the brain response in an experiment extends significantly into the pre-stimulus of the following trial, using the pre-stimulus period as reference can lead to the response of interest being suppressed. Such a suppression of brain responses might be the reason for the signal drops observed with SSP_data_ at n_PC_ > 4 and n_PC_ > 6 for single- and dual-axis data without dental wire, respectively. The first principal components may have been dominated by interference, but following ones included late components of the auditory response. In such a case, using suitable resting state data as reference instead may be advisable. Here, we unfortunately did not record resting state.

> *Recommendation 6:* When planning to use SSP on a participant with dental wire, record resting state as reference data.

We consciously used a noise covariance calculated from empty room data for the distributed source reconstruction to highlight the effect of the algorithms. In first tests using a noise covariance matrix computed from pre-stimulus data, even the unprocessed data with dental artifact achieved good source reconstruction, with a clear M100 activation in the right superior temporal region. This indicates that the inverse method in itself provides decent interference suppression and may make additional algorithms superfluous if one is only analysing activity at the source level. Other distributed source reconstruction methods and beamformers using such a noise covariance matrix would likely exhibit similar interference suppression, although this remains to be tested. As with SSP_data_, one should keep in mind that a clean pre-stimulus period is crucial. Otherwise, activation bleeding into the pre-stimulus period of the following trials can be suppressed.

This work started in response to users at NatMEG asking for guidance which methods to use to process their OPM-MEG data. As such, the investigation and results are limited to the HEDSCAN OPM-MEG system used locally and include only a limited number of algorithms that we considered most promising. OPM-MEG is very heterogenous, and systems come in a variety of shapes and sizes. The results may look different for another system. We hope that for readers using other OPM-MEG systems this work and related scripts can serve as an example and guide for how to evaluate and compare different interference suppression algorithms for their specific setup.

## Declaration of the use of AI

We did not use AI in writing this paper.

## Data and Code Availability

The analysis code and experimental paradigm used in this study is shared on GitHub: https://github.com/chrisNatMEG/opm_noise_algorithms.

The data is available at KI Data Repository: *[link to be added]*

## Author Contributions

C.P.: Conceptualization; Data curation; Formal analysis; Project administration; Software; Visualization; and Writing—original draft. D.L.: Conceptualization; Methodology; Resources; Supervision; and Writing—review & editing.

